# A machine-learning approach for detection of local brain networks and marginally weak signals identifies novel AD/MCI differentiating connectomic neuroimaging biomarkers

**DOI:** 10.1101/2021.07.29.454368

**Authors:** Yanming Li, Jian Kang, Chong Wu, Ivo D. Dinov, Jinxiang Hu, Prabhakar Chalise, Jonathan D. Mahnken, for the Alzheimer’s Disease Neuroimaging Initiative

## Abstract

**Introduction:** A computationally fast machine learning method is introduced for uncovering the wholebrain voxel-level connectomic spectra that differentiates different status of Alzheimer’s disease (AD). The method is applied to the Alzheimer’s Disease Neuroimaging Initiative (ADNI) Fluorine-fluorodeoxyglucose Positron Emission Tomography (FDG-PET) imaging and clinical data and identified novel AD/MCI differentiating connectomic neuroimaging biomarkers.

**Methods:** A divide-and-conquer algorithm is introduced for detect informative local brain networks at voxel level and whole-brain scale. The connection information within the local networks is integrated into the node voxels, which makes detection of the marginally weak signals possible. Prediction accuracy is significantly improved by incorporating the local brain networks and marginally weak signals.

**Results:** Brain connectomic structures differentiating AD and mild cognitive impairment (MCI), AD and healthy, and MIC and healthy were discovered. We identified novel AD/MCI-associated neuroimaging biomarkers by integrating local brain networks and marginally weak signals. For example, networkbased signals in paracentral lobule (p-value=6.1e-5), olfactory cortex (p-value=4.6e-5), caudate nucleus (1.8e-3) and precentral gyrus (1.8e-3) are informative in differentiating AD and MCI. Connections between calcarine sulcus and lingual gyrus (p-value=0.049), between parahippocampal gyrus and Amygdala (p-value=0.025), between rolandic opercula and insula lobes (p-values=0.0028 and 0.0026). An overall prediction accuracy of 95.3% was achieved by integrating the selected local brain networks and marginally weak signals, compared to 84.0% by not considering the inter-voxel connections and using marginally strong signals only.

**Conclusion:** (i) The connectomic structures differentiating AD and MCI are significantly different to that differentiating MCI and healthy, which may indicate different neuronal etiology for AD and MCI. (ii) Many neuroimaging biomarkers exert their effects on the outcome diseases through their connections to other markers. Integrating such connections can help identify novel neuroimaging biomarkers and improve disease prediction accuracy.

## 1. Introduction

Alzheimer’s disease (AD) ranks the third as a cause of death for people 75 and older^1,2^. Every 65 seconds, someone in the US develops AD^3^ and it afflicted around 5.8 million Americans in 2020^4^. Overall, AD is considered the most expensive disease^5^ in US, with the overall cost of the disease being $305 billion in the U.S. in the year 2020^4^. Before the onset of AD, individuals may experience an intermediate cognitive deterioration known as mild cognitive impairment (MCI) that are not severe enough to interfere with their daily activities. As the earliest detectable clinical stage toward AD, MCI provides an attractive checkpoint of disease-modifying intervention^6,7^. Compared to AD, MCI is more subtle and difficult to prognosis^8,9^. Early and accurate prediction of MCI and differentiating neuropathology of MCI and that of AD is crucially important for successful treatment development and precision prevention^10,11,12,13^.

Many cognitive tests have been developed for AD diagnosis^14^. However, these tests have certain limitations. For example, cognitive tests are often powerless in discriminating subtle differences between AD and MCI and between MCI and healthy. Also, cognitive tests cannot be used for novel neuroimaging biomarker discovery that may provide insights on underlying neuropathology.

High-resolution neuroimaging scans nowadays provide unparallel precision for discovering AD (MCI) associated neuroimaging biomarkers. For example, Positron Emission Tomography (PET) images^15,16,17^ have been successfully used to understand the neurodegenerative mechanisms^18^. Most of current methods focus on identifying the locales and individual values of imaging biomarkers, such as important voxels, hotspots or brain regions associated with a disease^19,20,21^, while ignoring the connectivity between markers. The number of potential connections between tens of thousands of voxels in a neuroimage is often of astronomical scales. For example, each ADNI PET scan used in our study consists of more than 185,000 voxels. Jointly inferring all potential connections between these voxels accounts to inverting a covariance matrix of dimension 185,000×185,000, which has a computational complexity of O(185, 000 ^2.375^)^22^. Even with very powerful computing tools, it is infeasible to invert such a large-dimensional matrix.

Even though extremely challenging, detection of voxel-level brain connectomes associated with AD has attracted much interest. First, both AD and MCI can be viewed as a connectomic disorder neuropathologically, in the sense that their onsite is often accompanied with not only the loss of brain matters itself, but also the reduction of inter-neuron fiber connectivity required for healthy cognitive functioning^23,24^. Brain connectomics, which models the whole brain as a spectrum of network circuits, provides a systematic view to AD neuropathology and have been increasingly used to link the diseases with structural and functional neuroimages^25^. Most existing brain connectomic networks are region-based functional networks, which aggregate neurons into priori-defined functionally related or spatially circumscribed regions of interest (ROIs)^26,27,28,29^. While the current neuropathological theories, such as beta-amyloid initiated senile plagues and tau-protein initiated neurofibrillary tangles, indicate that developments of AD and MCI are more directly related to breaks-down of connectivity at neuronal level, other than at regional level. Such connectomic patterns are more reflected in the voxel-level connections^30,31^. Region-based methods often lead to a loss of information and inferior prediction performance.

Another advantage of detecting inter-voxel connections is about prediction. As we will demonstrate shortly, a large portion of the predictiveness for AD is embedded in the connections between voxels. In fact, AD-risk imaging biomarkers currently identified explain only a small proportion of the disease variation^32,33,34^. More novel neuroimaging biomarkers are yet to be discovered. Integrating connections between voxels into the disease prediction makes the detection of marginally weak signals possible. The marginally weak signals have small power in differentiating the disease status and are not detectable by themselves. They are, therefore, usually ignored in contemporary neuroimaging association studies. However, when taken into consideration their connections with other signals, marginally weak signals could exude strong predictive effects^35,36^.

Such a case is illustrated in Figure 1. The left panel depicts a local brain-network in cerebellum crus on the left hemisphere consisting of twelve voxels. The network contains two marginally detectable voxels “vox 94316” and “vox 98031”. All other ten voxels are marginally weak and undetectable. The right-panel table in Figure 1 gives the marginal and local-network-adjusted mean differences between the AD and MCI groups. The first column lists the marginal mean differences (divided by the marginal standard deviations) without considering the connections between the voxels. The second column lists the localnetwork-adjusted mean differences. The local-network-adjusted mean difference for a voxel integrates its connective information with other voxels, such as the number of edges connected to it, the strength of these connections and marginal differentiating power of its connected voxels. The discriminant power of most of the ten marginally undetectable signals are significantly boosted up by incorporating their connective information. All the ten marginally weak voxels become more powerful in differentiating AD and MCI than the two marginally strong signals after adjusting for the local network structure.

**Figure 1:**
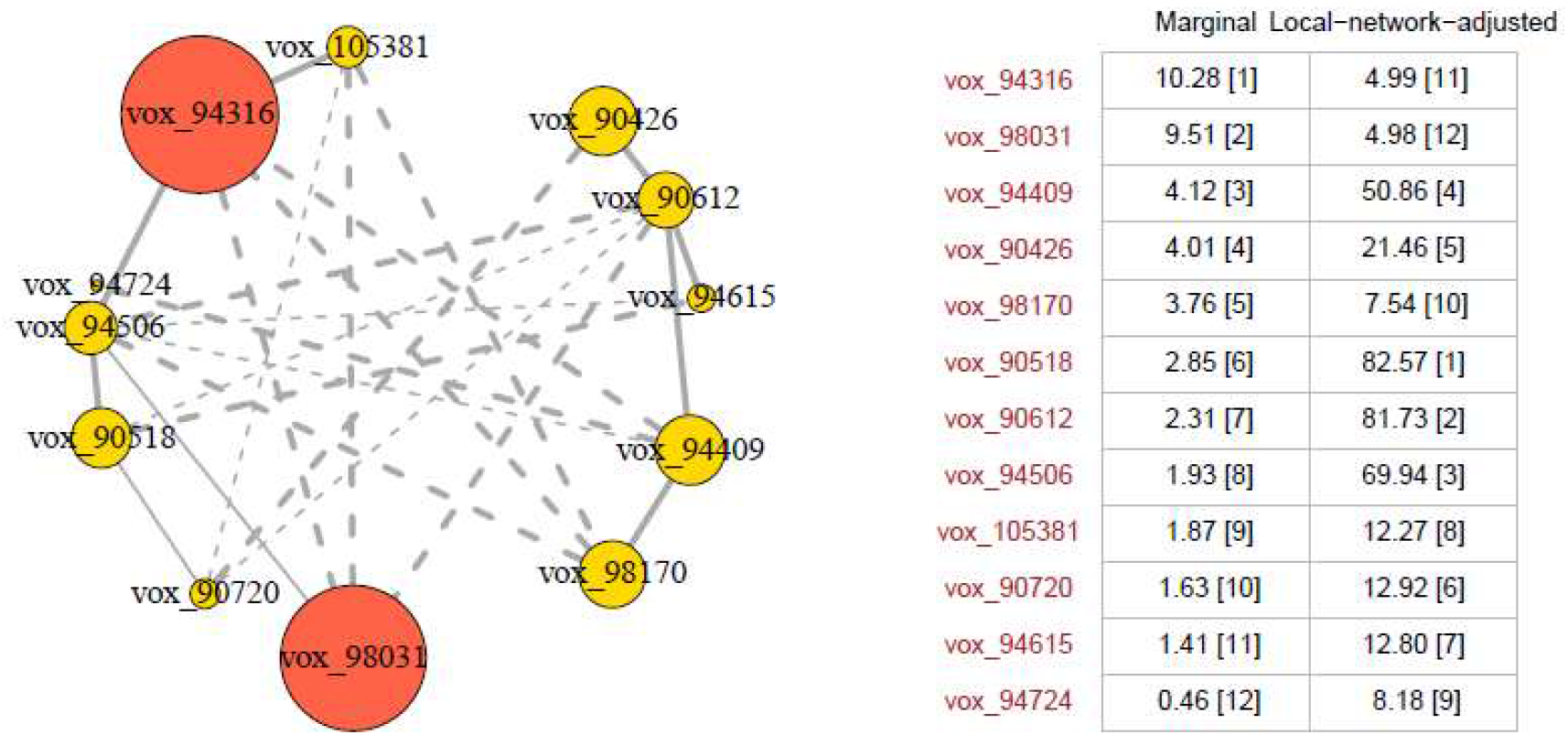
Left panel: a local brain-network in cerebellum Crus on the left hemisphere of the brain. Red: marginally detectable (strong) signal. Yellow: marginally undetectable (weak) signals. Solid line: positive connections. Dashed line: negative connections. Line width represents strength of connections. Right panel: Mean differences between the AD and MCI groups. First column: marginal mean differences (divided by the marginal variances). Second column: local network adjusted mean differences. Integers in the brackets are the ranks of the absolute mean differences for the 12 voxels, ranked from the greatest to the smallest.

In this work, we analyzed the ADNI Fluorine-fluorodeoxyglucose (FDG)-PET imaging and clinical data by detecting local brain networks and marginally weak signals^36^. To the best of our knowledge, this is the first whole-brain voxel-level connectivity study in the literature. The identified connectomic signatures were then integrated into AD/MCI prediction. Our approach avoids ultrahigh-dimensional precision matrix calculation by disassembling the whole brain connectome into disjoint local brain networks. It is highly efficient in computation. By integrating the voxel-level connectivity and marginally weak signals, the prediction accuracy has been significantly improved. Moreover, meaningful biological interpretation about identified network-based signatures can be made based on our findings, which might help to advance our understanding on the mechanisms of MCI and AD.

The rest of the paper is organized as following. Section 2 introduces the methods for detection of predictive local brain networks and marginally weak signals, and prediction rules for classifying the diseases. Section 3 introduces the ADNI PET imaging datasets. Section 4 gives the analysis results and biological annotations for our findings. The paper is concluded by Section 5, where a brief discussion of relevant issues is provided.

## 2. Methods

Figure 2 depicts the major steps of our analysis pipeline. Details for the methods used are elaborated next.

**Figure 2.**
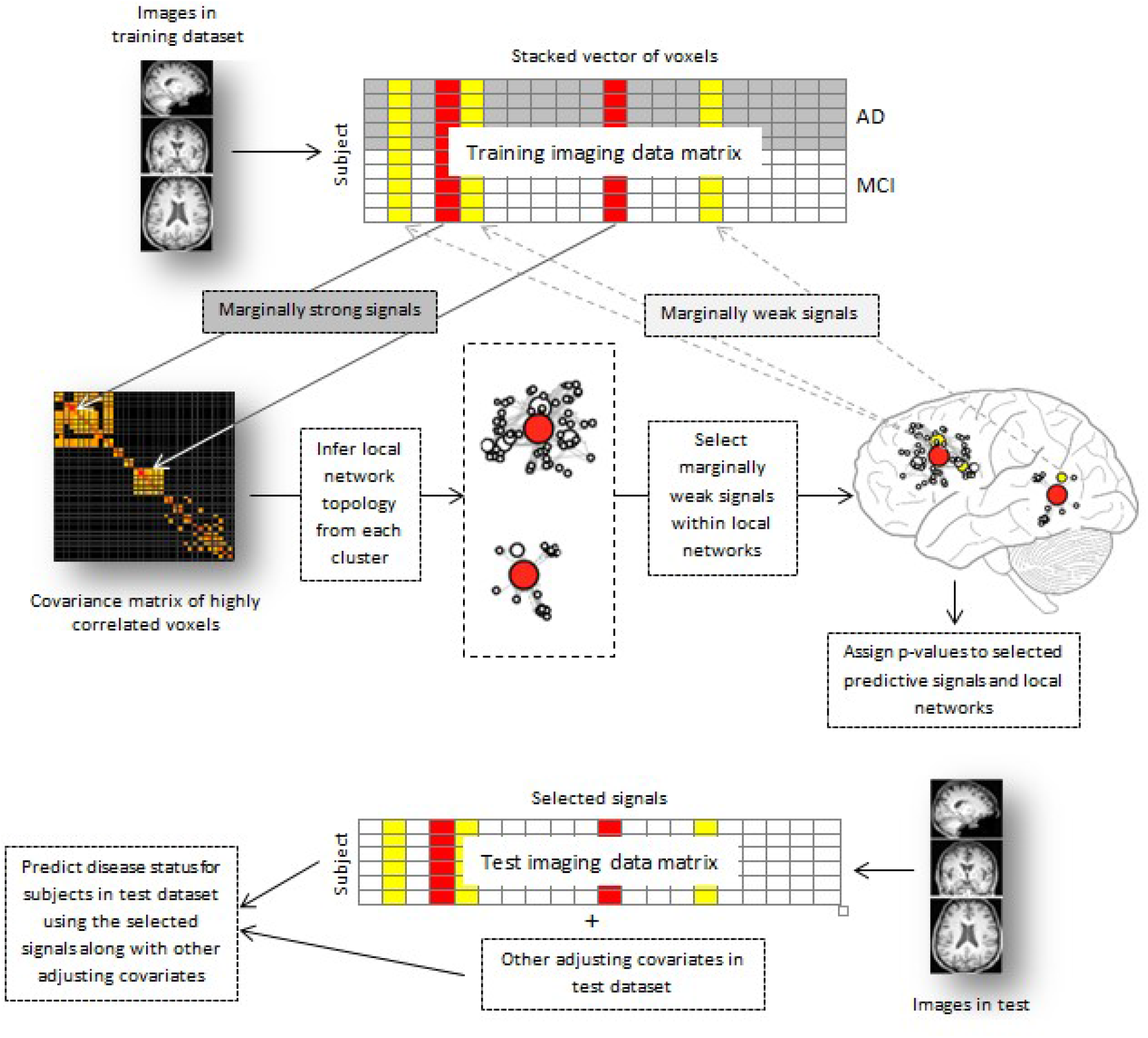
Flow chart of the proposed analysis pipeline.

### 2.1. Notations

Let *X* = (*X*_1_,…,*X_p_*)′ be the stacked vector for the intensity scores of all *p* voxels in an imaging scan. Denote by *X_i_* = (*X*_*i*,1_,…,*X_i,p_*)′ the observed *X* vector from subject *i*. Let *Y_i_* be the class indicator (coded as 0, 1, 2 for healthy, MCI and AD, respectively) for subject *i, i* = 1,…,*n*, where *n* is the total number of subjects. Denote by 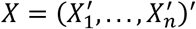 the *n* × *p* data matrix. Denote by *n_k_* the size of class k, k = 0, 1, 2. Denote by *G* = (*V, E*) the whole brain connectomic network, with a vertex set *V* ≡ {1,…,*p*} and an edge set *E*. A connection (or an edge) (*j, j*_0_) exists between two voxels *j* and *j*_0_ if and only if voxels *X_j_* and *X*_*j*_0__ are conditionally dependent given all other voxels. Note that connection (or an edge) between two voxels is essentially different than the correlation between them. The former is a joint concept depending on all the other voxels, while the latter is a marginal concept depending only on the pair of voxels under consideration.

### 2.2. Detection of marginally strong signals

Marginal two sample t-tests are used to select marginally strong signals, which each differentiates a pair of classes by itself. Specifically, for each feature *j, j* = 1,…, *p*, we calculate

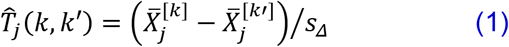

for a class pair (*k, k*′), *k* ≠ *k*′, in {0,1,2}. Here 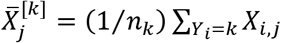 and 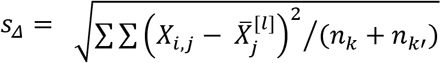. The first *τ* ≤ *n* features with the highest 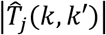 values are selected as the marginally strong signals. Here *τ* is a tuning parameter and can be selected by data-driven procedures such as cross-validation.

### 2.3. Detection of predictive local brain networks

Under the assumption that the *p* voxels follow a multivariate normal distribution with a mean vector *μ* and a covariance matrix *Σ*, The network *G* can be characterized by the precision matrix *Ω* = *Σ*^-1^. An edge (*j, j*′) exists if and only if the (*j, j*′)th entry of *Ω*, denoted by *Ω_jj′_*, is nonzero and the strength of the edge equals to the magnitude of *Ω_jj_*. As stated before, jointly estimating the whole *Ω* matrix of all 185,000 voxels is computationally prohibitive. Here we employed a “divide-and-conquer” algorithm introduced in Li et al.^36^, to detect the local brain networks. The local networks are much smaller in size and thus their corresponding precision matrices are much easier to calculate. Assume that each local brain network contains at least one marginally strong voxel. These marginally strong voxels serve as hubs in the local networks. For each marginally strong signal detected, we look for the set of voxels connected to it, either directly through an edge or indirectly through a path consisting of a series of edges. That is, to find the connected component in *Ω* containing it. However, it is impossible to detect such connected components without knowing *Ω*. Utilizing a statistical property which states that the connected component structure of *Ω* can be asymptotically recovered by that of the thresholded sample correlation matrix, 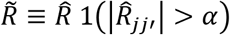)^37,38^, we can detect the connected components in 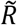 instead. Here 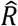 is the estimated sample correlation matrix and 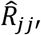 is its (*j, j*′)th entry and 1 is the indicator function. The thresholding parameter 0 < *α* < 1 controls the sparsity of 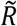. The computational complexity of 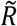 is orders of magnitude lighter than that of estimating *Ω*. Moreover, since the correlations can be estimated pairwise, they can be calculated in a parallel way on multi-core computer clusters.

Recursive labeling algorithm^39^ is employed in detecting the connected component in 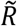 for each marginally strong signal. Denote by *C_l_*, *l* = 1,…,*B*, the *B* connected components identified. Each *C_l_* indexes a local brain network. Let 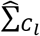 be the sub-sample covariance matrix corresponding to the set *C_l_*. The precision matrices 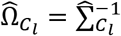 characterize the topology for the corresponding local networks. Sizes of *C_l_* s can be controlled by carefully choosing the thresholding parameter *α*.

### 2.4. Detection of marginally weak signals

Local-network-adjusted effects are then calculated for voxels within each *C_l_*. Specifically, for each *C_l_*, the following network-adjusted statistics vector is calculated:

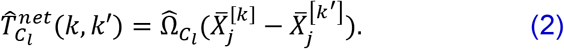

For each voxel *j* in *C_l_*, its network-adjusted statistic is the corresponding entry in 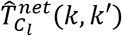. Specifically,

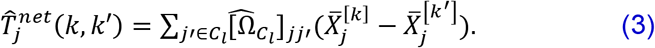

Compared to the marginal statistics 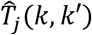 in (1), instead of standardizing by the marginal variation (measured in *s_Δ_*), 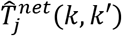 also adjusts for the local network connections for feature *j* (estimated in 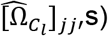) and the marginal differential effects of voxels connected to feature *j* (measured in 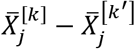 for *j*′ ≠ *j* in *C_l_*).

Not all the voxels in *C_l_*s are necessarily predictive. To reduce false positives, we further select predictive marginally weak signals within 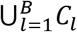. The top ranked *v* voxels with the greatest 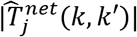 are selected, beside of the marginally strong signals. Here *v* is a tuning parameter controlling the size of marginally weak signal set. The predictive signals (both marginally strong and marginally weak), together with the local brain networks they are residing in, form the AD (or MCI) predictive brain connectome. Figure 3 gives a toy example about incorporating a local brain network into marginally weak signal detection.

**Figure 3.**
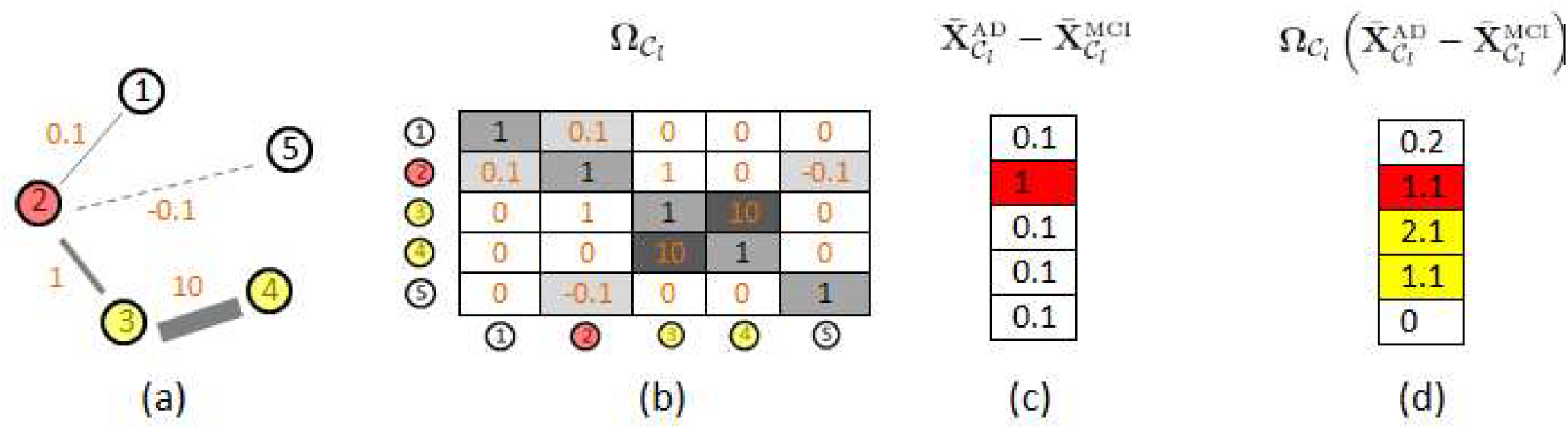
A toy example of a local brain network with five voxels. (a) Topology of the network. numbers on each edge are for the edge strength. Negative numbers mean the two voxels are negatively connected. (b) The corresponding precision matrix *Ω_C_l__*. (c) Marginal differentiating effects between AD and MCI groups. Red: marginally strong signals. (d) Local network adjusted differentiating effects between AD and MCI groups. Yellow: marginally weak signals.

### 2.5. Assigning p-values to selected signals and local brain networks

A non-parametric permutation test is used to evaluate significance of selected signals^40,41^. Specifically, the p-value for a selected voxel *j* is calculated through the following procedure:

1. Calculate 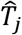 and 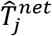 using the original data. If voxel *j* was not selected, then 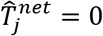.
2. Randomly permute the class label vector (*Y*_1_,…,*Y_n_*)^′^ *M* times for some large number *M* and generate *M* permuted datasets.
3. For each feature *j*, calculate (2) using the *m*th permuted dataset, denoted by 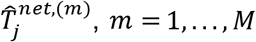.
4. Assign p-value to a selected voxel *j* to be

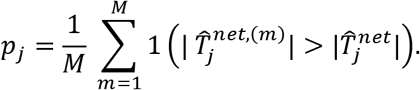

To access significance of the network-based signatures identified, we assign network p-values to the selected local brain networks using the following Hotelling’s T-squared distribution

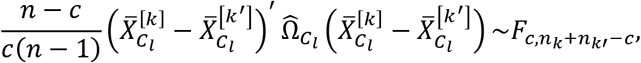

where, *c* is the number of voxels in *C_l_* and *F_c,n_k_+n_k′_−c_* is the F-distribution with parameters *c* and *n_k_* + *n_k′_ −c*.

### 2.6 Disease status prediction

Let *X_new_* and *Z_new_* be the predictor vector and adjusting covariate vector for a new subject in the test dataset, respectively, then the class of the new subject can be predicted by the following rule.

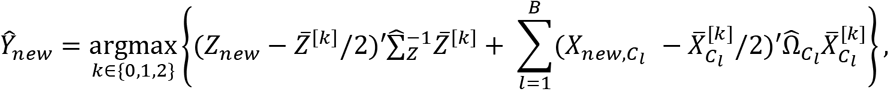

where *X_new,Cl_* is the subvector of *X_new_* indexed by *C_l_*.

## 3. ADNI datasets

During the last two decades, large amount of neuroimaging, genetic and clinical data have been acquired in ADNI studies^42,43,44^. In this study, we used the ADNI phase-I FDG-PET baseline scans with 54 AD patients, 131 MCI patients and 72 healthy subjects to detect brain connectomic structures for AD and MCI prediction.

### 3.1 Data processing

Standard preprocessing steps including co-registration, normalization and spatial smoothing (8 mm full width at half maximum) were applied to the PET dataset. Each image is registered to a template with 185,405 active voxels embedded in a 91 x 109 x 91 3D-array. We further grouped the 185,405 voxels into 116 regions of interest (ROI) segmented by the automated anatomical labeling atlas^45^. ROI names and numbers are listed in the Table A1 in the Appendix.

## 4. Analysis and results

Pairwise correlations and covariances between all voxels were first calculated on parallel threads using a Hewlett-Packard workstation with 16 core Intel Xeon processors and a 256GB memory size. Correlation matrix was then thresholded at 0.8 to retain only highly correlated voxels. To further reduce the computational burden, we set all entries within a thresholded correlation block between two ROIs to zeros if the average *L*_2_ norm (*L*_2_ norm divided by the number of nonzero entries in the block) of that block is ≤ 0.4. Thus, only highly correlated ROIs were be considered. For example, For ROI 5, only its closely connected ROIs: ROI 6, 9, 15, 25, 26, 27 and 28, were considered in our analysis (see Figure 4-a). This correlation block structure represents an overarching region-level connectivity structure (Figure 4-b).

**Figure 4.**
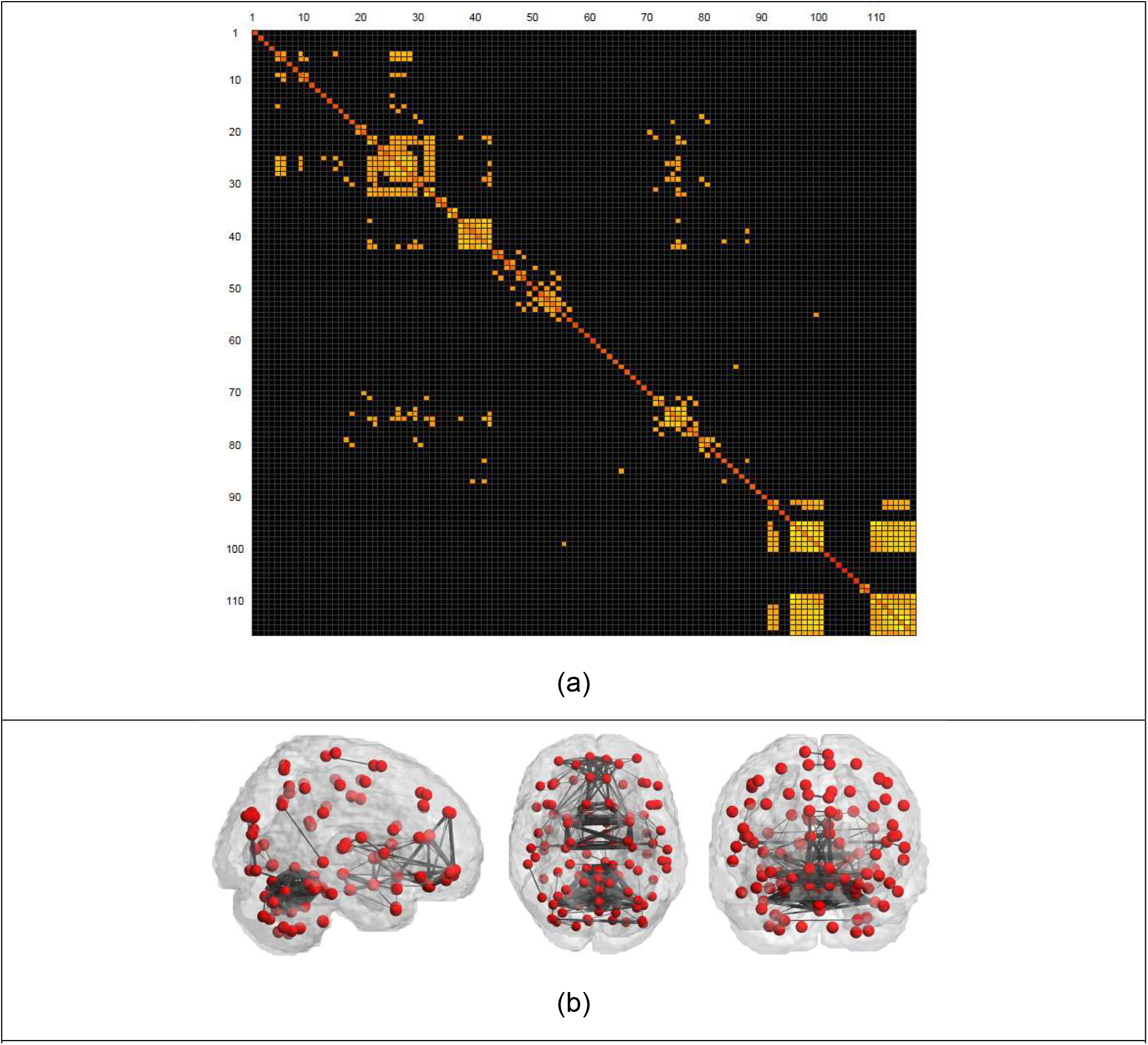
(a) ROI block-wise thresholded sample correlation matrix. A cell represents the weighted *L*_2_ norm of the correlation block between a pair of ROI’s (weighted by 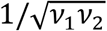 where *v*_1_ and *v*_2_ are the numbers of voxels in the two ROIs, respectively) in the voxel-level correlation matrix. Raw voxel-level correlations are thresholded at 0.8. The weighted *L*_2_ norms are further thresholded at 0.8, which is the median of the 6,670 off-diagonal ROI block *L*_2_ norms. All blocks with *L*_2_ norm ≤ 0.4 are set to zero. The black squares represent the zero blocks. The lighted squares are for non-zero blocks. Lighter color indicates greater *L*_2_ norm values. (b) 3D ROI connection topology viewed from sagittal (left), axial (middle) and coronal (right) directions. Each dot represents a central point of an ROI. An edge exists between two ROIs if the corresponding thresholded correlation block is nonzero.

Variable selection was performed separately within each highly correlated ROI clusters. Parallel computing was implemented for this step. We used *τ* = 5, *α* = 0.8 and *v* = 20 on each ROI cluster to select local-brain networks, and marginally strong and weak signals that differentiated a pair of classes. When searching for the connected component of a marginally strong signal, we focused on the voxels that are connected to a marginally strong signal with a connection path less than 10 in length. We used *M* = 10,000 in the non-parametric permutation tests. In total, 2,686 informative voxels were selected. Among them, 836 are marginally strong and 1,850 are marginally weak. There were 661 (302), 695 (306) and 610 (309) were marginally weak (marginally strong) signals for differentiating AD and MCI, AD and Healthy, and MCI and Healthy, respectively. There were 47 (61) marginally weak (marginally strong) signals that differentiate both AD-to-MCI and AD-to Healthy, 3 (0) marginally weak (marginally strong) signals that differentiate both AD-to-MCI and MCI-to-Healthy, 67 (20) marginally weak (marginally strong) signals that differentiate both AD-to Healthy and MCI-to-Healthy, and 1 marginally weak signal that differentiate all three pairs. The selected informative signals differentiating AD and MCI, AD and healthy, and MCI and healthy are sitting in 168, 129 and 122 local brain networks, respectively. Table 1 lists the top selected voxel signals that are informative in differentiating at least one pair of disease status. Figure 5 shows the top selected marginally weak signals, along with their marginal and network-adjusted effects. Note that some regions identified containing mostly marginally weak signals have either not been reported to be associated with AD or MCI, or just been discovered recently, such as middle temporal gyrus^46^, olfactory bulbs^47^, lingual gyrus^48^ and amygdala^69^. These novel findings demonstrate the power of our method in identifying novel neuroimaging biomarkers. Table 2 lists the top selected brain networks with a significant unadjusted p-values. Most of selected voxels would still be significant after adjusting for multiple tests. The topologies of these networks are depicted in Figure 6. For the sake of presentation, we only included 12 the most closely connected voxels to a marginally strong signal in each depicted network. The whole brain connectomic structure that differentiate AD and MCI is presented in Figures 7 and 8. Each cluster of arcs in Figure 7 represent connections within an ROI. Arcs run across different clusters represent connections between different ROIs. Figure 8 shows, on the other hand, the positions and overarching topologies of the detected connectome. The corresponding figures for connectomes that differentiate AD and healthy, MCI and healthy are provided in the Appendix.

**Figure 5:**
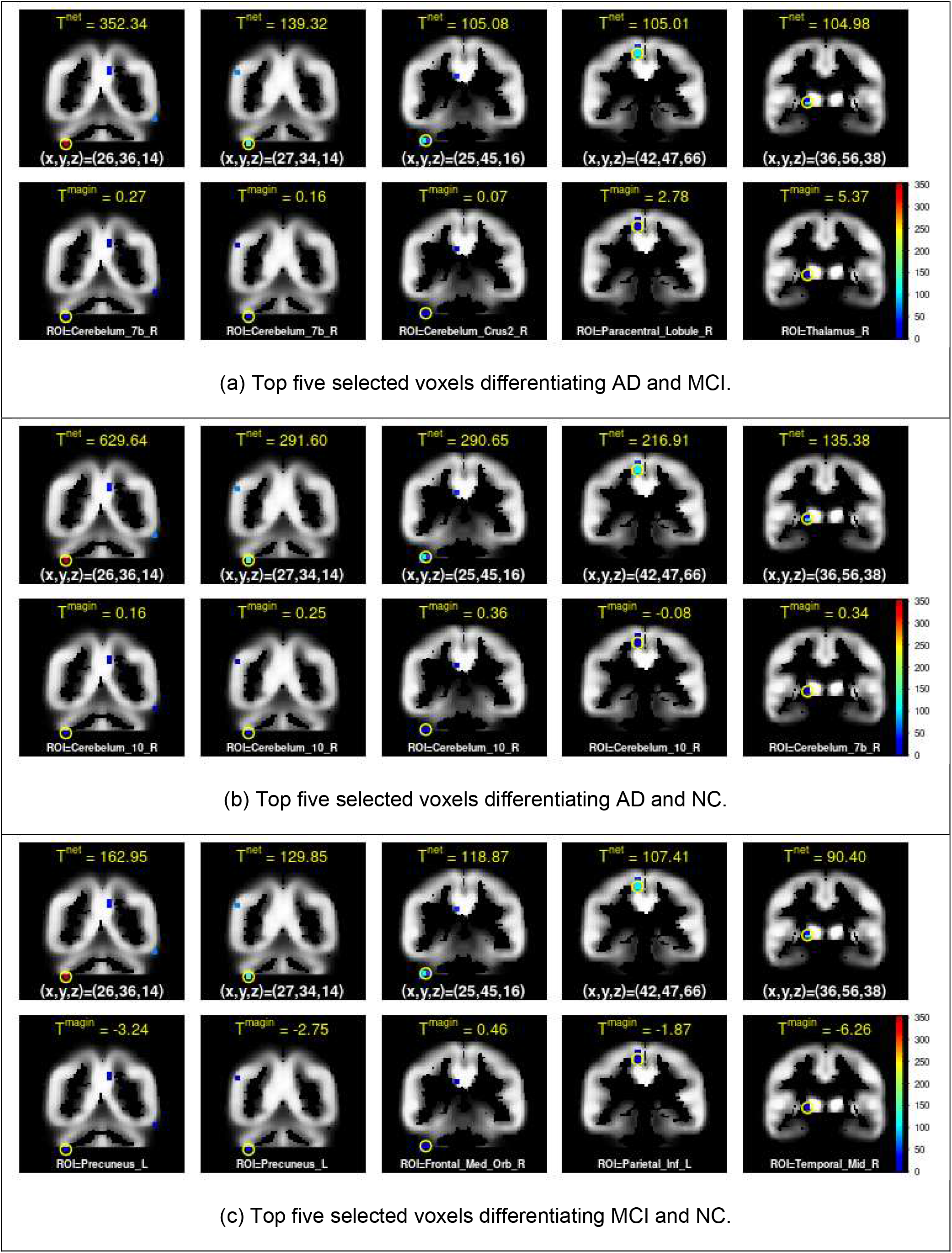
The symbol 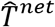, defined in equation (3), represents the local network adjusted mean difference between a pair of classes for a voxel. The symbol 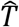, defined in equation (1), represents the marginal mean difference between a pair of classes for a voxel. The triplet (x, y, z) represents the coordinates of the top voxel signal highlighted in yellow circle. ROI gives the regions of interest for the top voxel signal.

**Figure 6:**
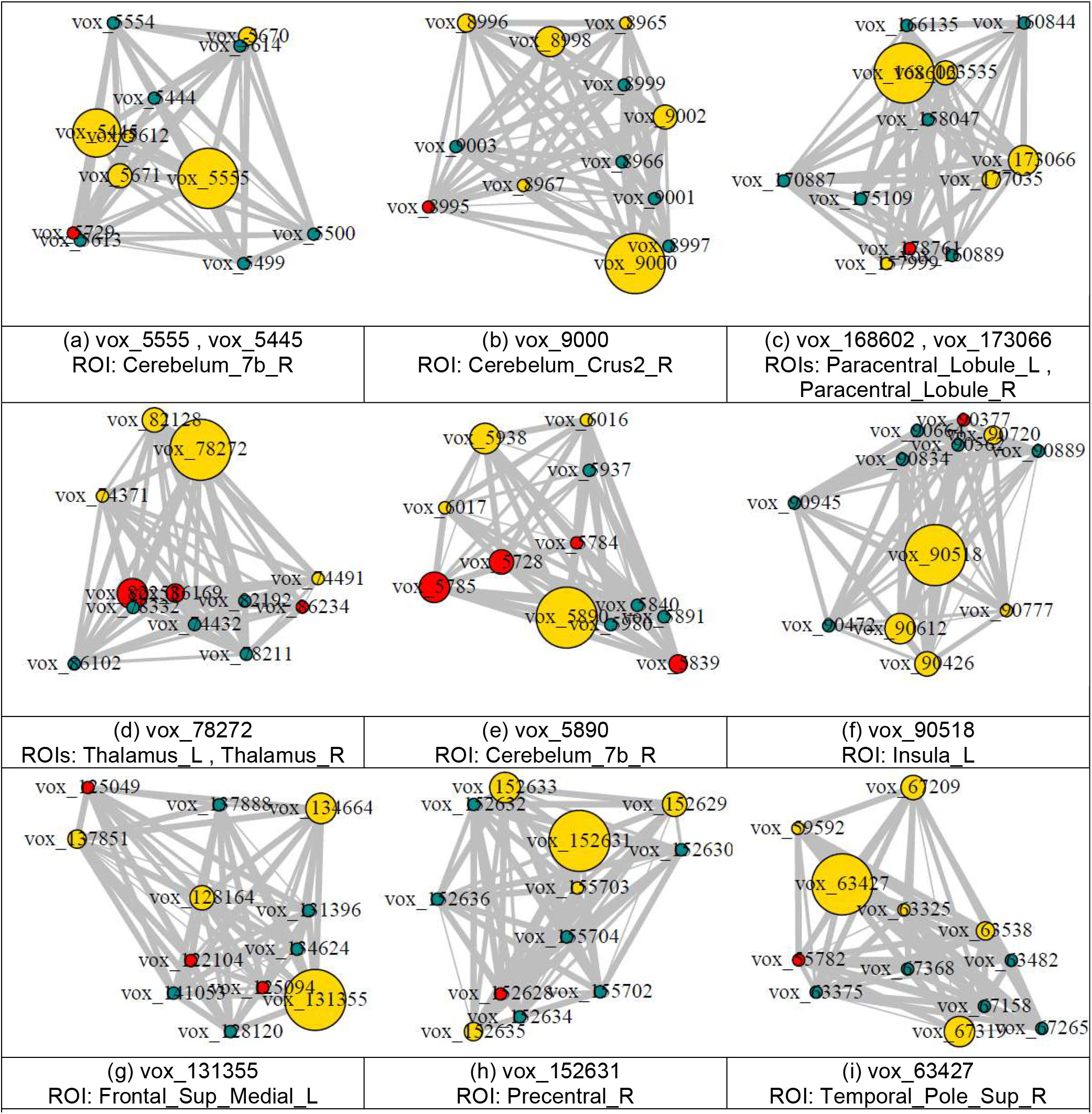
Example local brain networks that differentiate AD and MCI groups. Yellow: marginally weak signals. Red: marginally strong signals. Blue: other selected connecting signals that differentiate other pairs. Node sizes are proportional to the local-network-adjusted mean differences. Edge widths are proportional to the absolute partial correlations. The top selected voxels within each network and the ROI(s) the network resided in are indexed.

**Figure 7:**
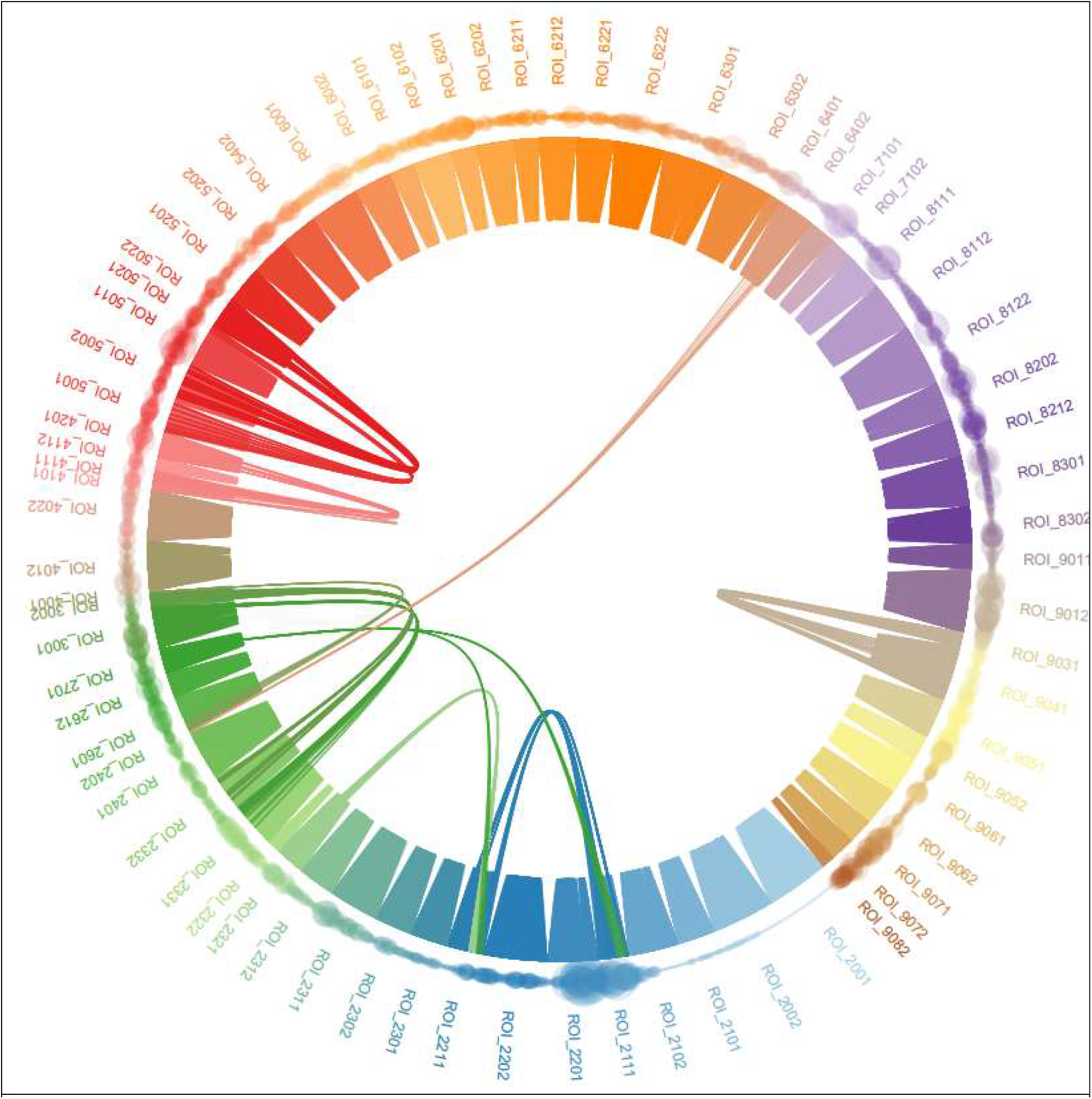
Local brain network connectome differentiating AD and MCI. Each dot represents a voxel and each cluster of arcs represents a local network detected. Different colors represent different ROIs. A network connecting dots in different colors indicates that the network is crossing multiple ROIs.

**Figure 8:**
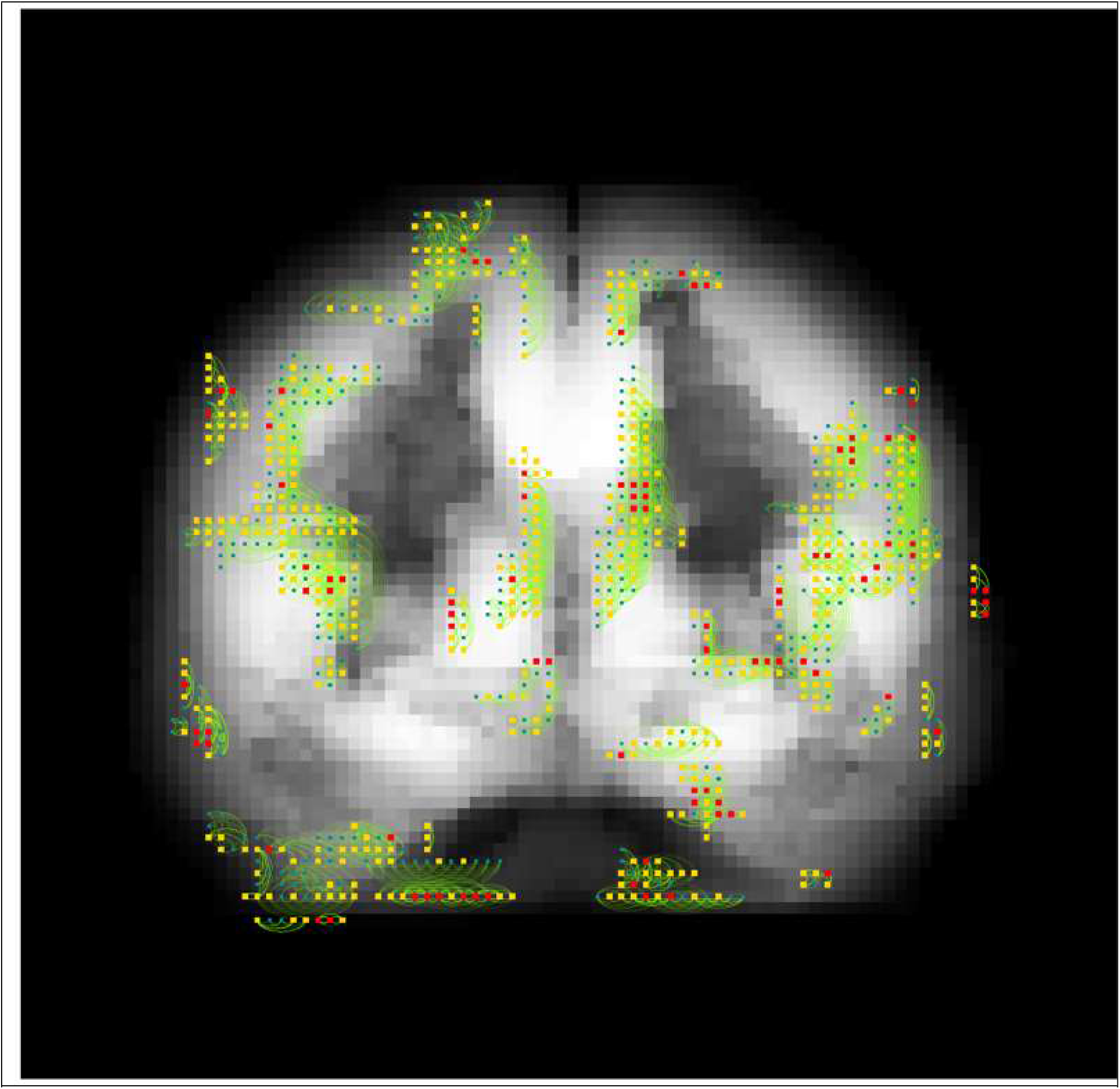
Overlaying of the brain network connectome differentiating AD and MCI on the brain. All local networks are projected to a coronal slice of the brain at midline. Red voxel: marginally strong signal. Yellow voxel: marginally weak signal. Blue voxel: non-selected connecting voxel in a selected local brain network (might differentiate other pair of classes).

**Table 1.**
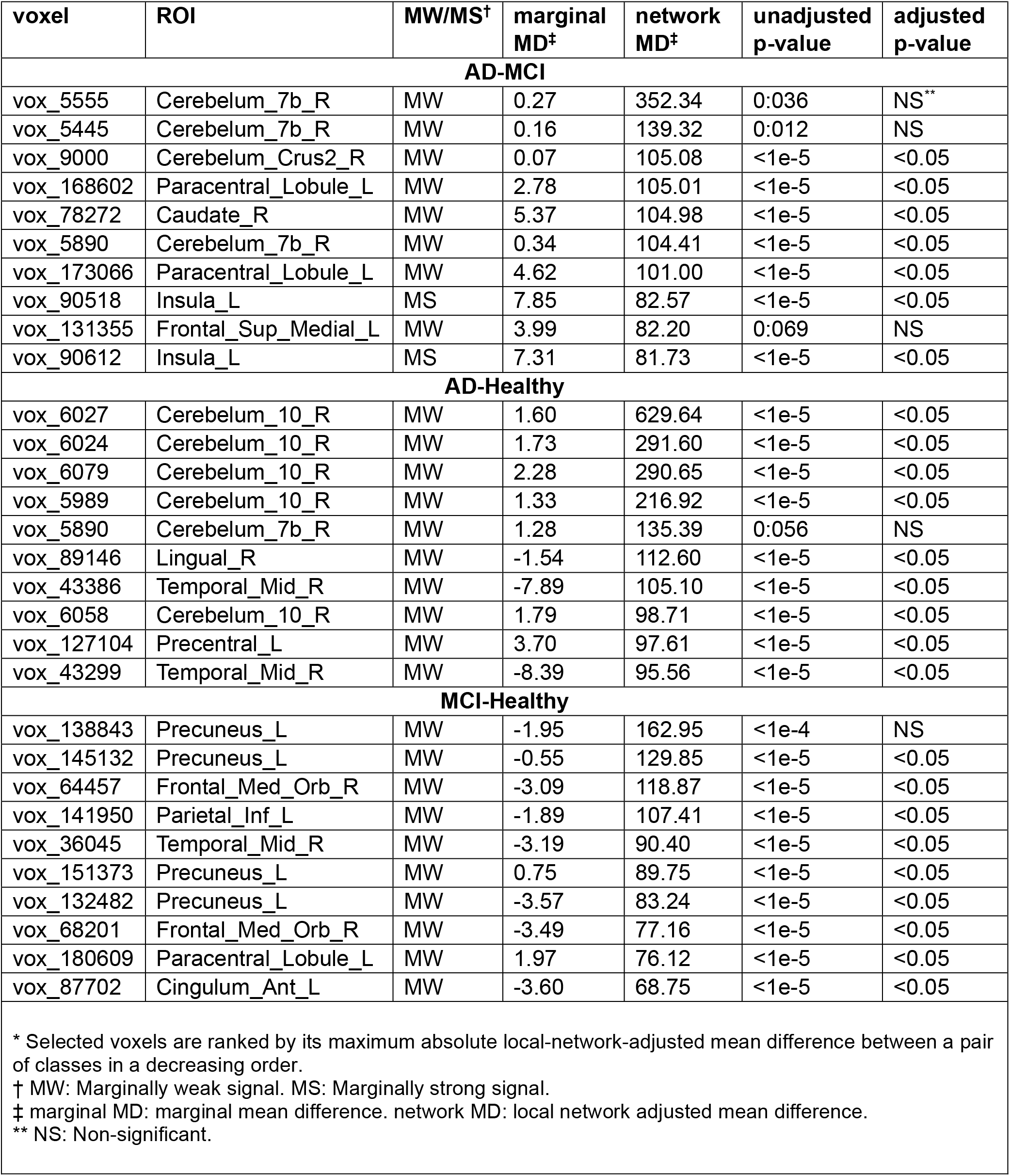
Top selected voxel signals*.

**Table 2:** Top selected local brain networks*.

| top voxel(s) | ROI(s) the network sitting in | MW<sup>†</sup><br># | MS<sup>‡</sup><br># | Hotelling<br>t-squared<br>statistics | unadjusted<br>network<br>p-value | Adjusted**<br>network<br>p-value | literature<br>evidence |
| --- | --- | --- | --- | --- | --- | --- | --- |
| **AD-MCI** |  |  |  |  |  |  |  |
| vox_168602 | Paracentral_Lobule_L | 5 | 1 | 5.7 | 6.1e-5 | 0.01 | 52,53,54,55 |
| vox_78272 | Caudate_R | 4 | 3 | 3.7 | 1.8e-3 | NS | 56,57,58 |
| vox_90518 | Olfactory_L | 5 | 1 | 5.9 | 4.6e-5 | 0.008 | 59,60,53,61 |
| vox_152631 | Precentral_R | 5 | 1 | 4.0 | 0.0019 | NS | 52,62,63 |
| **AD-Healthy** |  |  |  |  |  |  |  |
| vox_6027;<br>vox_5989;<br>vox_6024;<br>vox_5987 | Cerebelum_10_L;<br>Cerebelum_10_R | 5 | 3 | 3.5 | 0.03 | NS | 64 |
| vox_89146 | Lingual_R;<br>Calcarine_R | 5 | 1 | 7.5 | 3.8e-6 | 0.0005 | 52,65,66,67,68 |
| vox_32891 | Amygdala_L | 24 | 1 | 4.7 | 1.6e-8 | 2.1e-6 | 69 |
| **MCI-Healthy** |  |  |  |  |  |  |  |
| vox_138843;<br>vox_145132;<br>vox_151373;<br>vox_132482 | Precuneus_L | 5 | 1 | 8.5 | 2.7e-7 | 3.5e-5 | 70,71,72,73 |
| vox_64457;<br>vox_68201 | Frontal_Med_Orb_R | 6 | 1 | 2.7 | 0.014 | NS | 52,74 |
| vox_141950 | Parietal_Inf_L | 4 | 3 | 2.3 | 0.038 | NS | 75,76 |
| vox_36045 | Temporal_Mid_R | 6 | 1 | 3.5 | 0.0016 | NS | 77,78 |
| * Local brain networks with a significant unadjusted network p-value and host selected informative voxels are listed.<br>** Adjusted p-values are adjusted for the number of local networks selected that differentiate the pair of disease classes.<br>† MW #: number of selected marginally weak signals in the network.<br>‡ MS #: number of selected marginally strong signals in the network. |  |  |  |  |  |  |  |

For many of the local networks identified, associations between their residing ROI and AD status are supported by literature evidence. These threads of evidence are listed in the last column of Table 2. Our analysis showed that connections within paracentral lobule (p-value=6.1e-5), olfactory cortex (p-value=4.6e-5), caudate nucleus (1.8e-3) and precentral gyrus (1.8e-3) are informative in differentiating AD and MCI; Connections within cerebellum crus (p-value=0.03), lingual gyrus and calcarine sulcus (p-value=3.8e-6) and amygdala (p-value=1.6e-8) are informative in differentiating AD and the healthy; Connections within precuneus gyrus (p-value=2.7e-7), middle temporal gyrus (p-value=1.6e-3), middle orbitofrontal cortex (p-value=0.014) and inferior parietal lobule (p-value=0.038) are informative in differentiating MCI and the healthy.

Connections across different ROIs that contribute to differentiating a pair of disease status are of particular interest, as they may indicate the functional connection that contributes to AD etiology. The following cross-ROI connections were identified for differentiating AD and MCI: connections between calcarine sulcus and lingual gyrus (network p-value=0.049), connections between parahippocampal gyrus and amygdala (network p-value=0.025), connections between rolandic opercula and insula lobes (network p-values=0.0028 in the left hemisphere and =0.0026 in the right hemisphere). The connections between calcarine sulcus and lingual gyrus (network p-value=7.5e-6) and connections between parahippocampal gyrus and amygdala (network p-value=1.7e-4), were also identified to differentiate AD and the healthy. Other significant cross-ROI networks that differentiate AD and the healthy, MCI and the healthy are listed in Table S2 in the Appendix.

The patterns of the selected connectomes differentiating the three disease pairs (AD-to-MCI, AD-to-healthy, and MCI-to-healthy) are significantly different. See Figure 8 and Figures A3 and A6 in the Appendix. While connections between calcarine sulcus and lingual gyrus, between parahippocampal gyrus and amygdala are informative for differentiating both AD-to-MCI and AD-to-healthy; connections between cingulum posterior, left hemisphere and cingulum posterior, right hemisphere, between cingulum middle, left hemisphere and cingulum middle, right hemisphere are informative for differentiating both AD-to-healthy and MCI-to-healthy. Note that there is no selected local network that differentiates both AD-to-MCI and MCI-to-healthy, which indicates that the development of MCI and the progression from MCI to AD may have different neuropathology pathways.

For prediction, we used five-fold cross-validation to divide the whole dataset into five parts with about the same sample sizes. Each time, we combined four folds into a training set and set the rest fold as the test set. We applied our analysis procedure on the training dataset to select predictive voxels and networks. Same tuning parameters *τ* = 5, *α* = 0.8 and *v* = 20 were used on each ROI cluster. The selected voxels on 116 ROIs were then pooled together and used to predict the disease status on the test dataset, along with covariates age and sex. We repeated this procedure till each fold has been used as the test set once. The overall classification errors were then computed by summing over all five test sets. The prediction results are summarized in Table 3. For comparison, we also reported prediction results from (i) marginal linear discriminant analysis (LDA), which assumes all voxels are independent (unconnected), (ii) sure independent screening (SIS)^49,50^, and (iii) iterative sure independent screening (ISIS)^50,51^. For SIS and ISIS, logistic regressions were applied on each pair of classes. The class with the highest average predicted probability was assigned to be the final predicted class. Numbers of misclassification are given in Table 3. Our approach gives the smallest prediction errors overall and within each class.

**Table 3:** Numbers of mis-classified subjects by different methods.

|  | **AD** | **MCI** | **Healthy** | **total** |
| --- | --- | --- | --- | --- |
| localNet-LDA | 2 | 7 | 3 | 12 |
| LDA | 5 | 28 | 8 | 41 |
| SIS-logistic | 11 | 46 | 26 | 83 |
| ISIS-logistic | 7 | 48 | 10 | 65 |

## 5. Discussion

The computational burden of the proposed analysis pipeline is comparable to that from the marginal approaches^49,50^. The major computational cost comes from calculating the whole brain correlation matrix. However, this can be alleviated by parallel computing. The connected component searching could also be computation-intensive if the number of connected voxels to a marginally strong signal is large. In our application, we restricted the length of any connection path in a connected component to be less than 10. This way, only the most closely connected voxels were maintained in a large-sized connected component. The analysis source codes were packed into an R package and can be found at https://github.com/lyqglyqg/mLDA

### Strengths and limitations

The strengths of this study are two-fold. First, it integrates inter-voxel connectivity into neuroimaging signal selection, and makes it possible to select both network-based and marginally weak signals. Secondly, different to many principal component analyses in neuroimaging association studies, our approach can select neuropathologically meaningful biomarkers - the local brain networks, while achieving high prediction accuracy.

Accurate estimation for the covariance structures requires greater sample sizes compared to for the mean structures. Breaking down the overall brain connectomic structures onto separate ROI clusters (see Figure 4) may cause loss of information. It is recognized that the inter-voxel connectivity may not fully reveal the connection pattern between neurons.

