## Supplemental Tables and Figures for "A machine-learning approach for detection of local brain networks and marginally weak signals identifies novel AD/MCI differentiating connectomic neuroimaging biomarkers"

### Appendix

**Table A1**: ROI ids and names.

| ROI id | ROI name | ROI id | ROI name | ROI id | ROI name |
|---|---|---|---|---|---|
| 2001 | Precentral_L | 4112 | ParaHippocampal_R | 8101 | Heschl_L |
| 2002 | Precentral_R | 4201 | Amygdala_L | 8102 | Heschl_R |
| 2101 | Frontal_Sup_L | 4202 | Amygdala_R | 8111 | Temporal_Sup_L |
| 2102 | Frontal_Sup_R | 5001 | Calcarine_L | 8112 | Temporal_Sup_R |
| 2111 | Frontal_Sup_Orb_L | 5002 | Calcarine_R | 8121 | Temporal_Pole_Sup_L |
| 2112 | Frontal_Sup_Orb_R | 5011 | Cuneus_L | 8122 | Temporal_Pole_Sup_R |
| 2201 | Frontal_Mid_L | 5012 | Cuneus_R | 8201 | Temporal_Mid_L |
| 2202 | Frontal_Mid_R | 5021 | Lingual_L | 8202 | Temporal_Mid_R |
| 2211 | Frontal_Mid_Orb_L | 5022 | Lingual_R | 8211 | Temporal_Pole_Mid_L |
| 2212 | Frontal_Mid_Orb_R | 5101 | Occipital_Sup_L | 8212 | Temporal_Pole_Mid_R |
| 2301 | Frontal_Inf_Oper_L | 5102 | Occipital_Sup_R | 8301 | Temporal_Inf_L |
| 2302 | Frontal_Inf_Oper_R | 5201 | Occipital_Mid_L | 8302 | Temporal_Inf_R |
| 2311 | Frontal_Inf_Tri_L | 5202 | Occipital_Mid_R | 9001 | Cerebelum_Crus1_L |
| 2312 | Frontal_Inf_Tri_R | 5301 | Occipital_Inf_L | 9002 | Cerebelum_Crus1_R |
| 2321 | Frontal_Inf_Orb_L | 5302 | Occipital_Inf_R | 9011 | Cerebelum_Crus2_L |
| 2322 | Frontal_Inf_Orb_R | 5401 | Fusiform_L | 9012 | Cerebelum_Crus2_R |
| 2331 | Rolandic_Oper_L | 5402 | Fusiform_R | 9021 | Cerebelum_3_L |
| 2332 | Rolandic_Oper_R | 6001 | Postcentral_L | 9022 | Cerebelum_3_R |
| 2401 | Supp_Motor_Area_L | 6002 | Postcentral_R | 9031 | Cerebelum_4_5_L |
| 2402 | Supp_Motor_Area_R | 6101 | Parietal_Sup_L | 9032 | Cerebelum_4_5_R |
| 2501 | Olfactory_L | 6102 | Parietal_Sup_R | 9041 | Cerebelum_6_L |
| 2502 | Olfactory_R | 6201 | Parietal_Inf_L | 9042 | Cerebelum_6_R |
| 2601 | Frontal_Sup_Medial_L | 6202 | Parietal_Inf_R | 9051 | Cerebelum_7b_L |
| 2602 | Frontal_Sup_Medial_R | 6211 | SupraMarginal_L | 9052 | Cerebelum_7b_R |
| 2611 | Frontal_Med_Orb_L | 6212 | SupraMarginal_R | 9061 | Cerebelum_8_L |
| 2612 | Frontal_Med_Orb_R | 6221 | Angular_L | 9062 | Cerebelum_8_R |
| 2701 | Rectus_L | 6222 | Angular_R | 9071 | Cerebelum_9_L |
| 2702 | Rectus_R | 6301 | Precuneus_L | 9072 | Cerebelum_9_R |
| 3001 | Insula_L | 6302 | Precuneus_R | 9081 | Cerebelum_10_L |
| 3002 | Insula_R | 6401 | Paracentral_Lobule_L | 9082 | Cerebelum_10_R |
| 4001 | Cingulum_Ant_L | 6402 | Paracentral_Lobule_R | 9100 | Vermis_1_2 |
| 4002 | Cingulum_Ant_R | 7001 | Caudate_L | 9110 | Vermis_3 |
| 4011 | Cingulum_Mid_L | 7002 | Caudate_R | 9120 | Vermis_4_5 |
| 4012 | Cingulum_Mid_R | 7011 | Putamen_L | 9130 | Vermis_6 |
| 4021 | Cingulum_Post_L | 7012 | Putamen_R | 9140 | Vermis_7 |
| 4022 | Cingulum_Post_R | 7021 | Pallidum_L | 9150 | Vermis_8 |
| 4101 | Hippocampus_L | 7022 | Pallidum_R | 9160 | Vermis_9 |
| 4102 | Hippocampus_R | 7101 | Thalamus_L | 9170 | Vermis_10 |
| 4111 | ParaHippocampal_L | 7102 | Thalamus_R | | |

- "_L": left hemisphere; "_R": right hemisphere; "_Sup": superior; "_Mid": middle; "_Inf": inferior.

**Table A2**: Selected crossing-ROI netwoks with significant marginal p-values.

| pair | ROI 1 | ROI 2 | p-value |
|---|---|---|---|
| AD-MCI | Calcarine_R<br>ROI_5002 | Lingual_R<br>ROI_5022 | 0.049 |
| | ParaHippocampal_L<br>ROI_4111 | Amygdala_L<br>ROI_4201 | 0.025 |
| | Rolandic_Oper_L<br>ROI_2331 | Insula_L<br>ROI_3001 | 0.0028 |
| | Rolandic_Oper_R<br>ROI_2332 | Insula_R<br>ROI_3002 | 0.0026 |
| AD-Healthy | Calcarine_R<br>ROI_5002 | Lingual_R<br>ROI_5022 | 7.5e-6 |
| | Calcarine_L<br>ROI_5001 | Lingual_L<br>ROI_5021 | 4.4e-6 |
| | Amygdala_L<br>ROI_4201 | Cingulum_Mid_L<br>ROI_4111 | 1.7e-4 |
| | Cingulum_Post_L<br>ROI_4021 | Cingulum_Post_R<br>ROI_4022 | 1.2e-9 |
| | Cingulum_Mid_L<br>ROI_4011 | Cingulum_Mid_R<br>ROI_4012 | 4.3e-6 |
| MCI-Healthy | Cingulum_Post_L<br>ROI_4021 | Cingulum_Post_R<br>ROI_4022 | 0.0021 |
| | Cingulum_Mid_L<br>ROI_4011 | Cingulum_Mid_R<br>ROI_4012 | 4.1e-4 |
| | Temporal_Pole_Mid_L<br>ROI_8211 | ParaHippocampal_L<br>ROI_4111 | 0.046 |

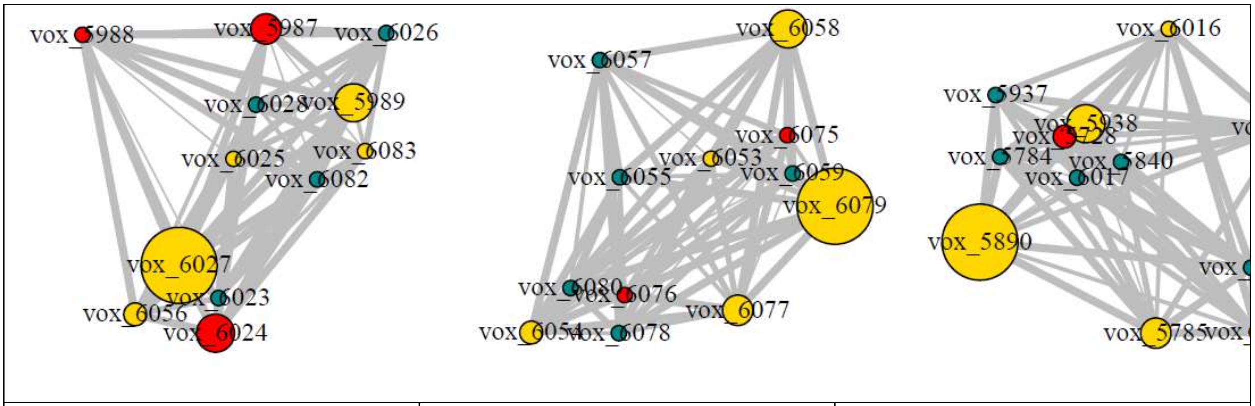


| (a) vox_6027 & vox_5989<br>vox_sss6024 & vox_5987<br>ROIs: Cerebelum_10_R &<br>Cerebelum_10_L | (b) vox_6079 & vox_6058<br>ROI: Cerebelum_10_R | (c) vox_5890<br>ROI: Cerebelum_7b_R |
|---|---|---|

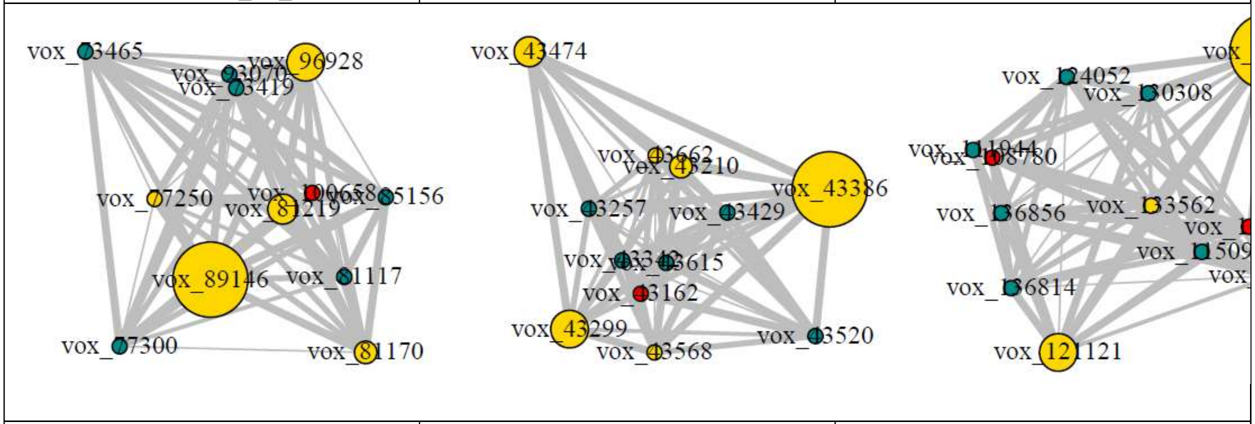


| (d) vox_89146<br>ROIs: Lingual_R & Calcarine_R | (e) vox_43386 & vox_43299 &<br>Vox_43474<br>ROI: Temporal_Mid_R | (f) vox_127104<br>ROI: Precentral_L |
|---|---|---|

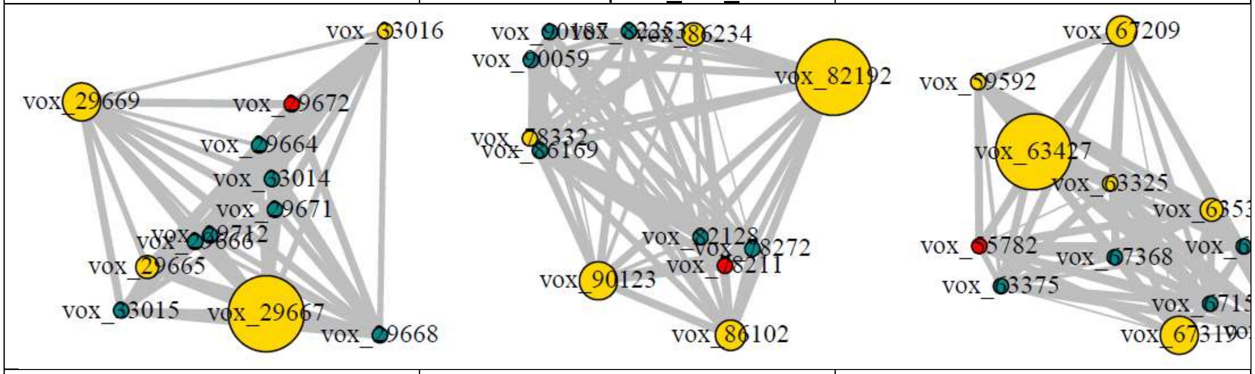


| (g) vox_29667<br>ROI: Temporal_Pole_Sup_R | (h) vox_82192<br>ROI: Thalamus_R | (i) vox_166077<br>ROI: Postcentral_R |
|---|---|---|

**Figure A1:** Example local brain networks that differentiate AD and healthy groups. Yellow: marginally weak signals. Red: marginally strong signals. Blue: other selected connecting signals that differentiate other pairs. Node sizes are proportional to the local-network-adjusted mean differences. Edge widths are proportional to the absolute partial correlations. The top selected voxels within each network and the ROI(s) the network resided in are indexed.

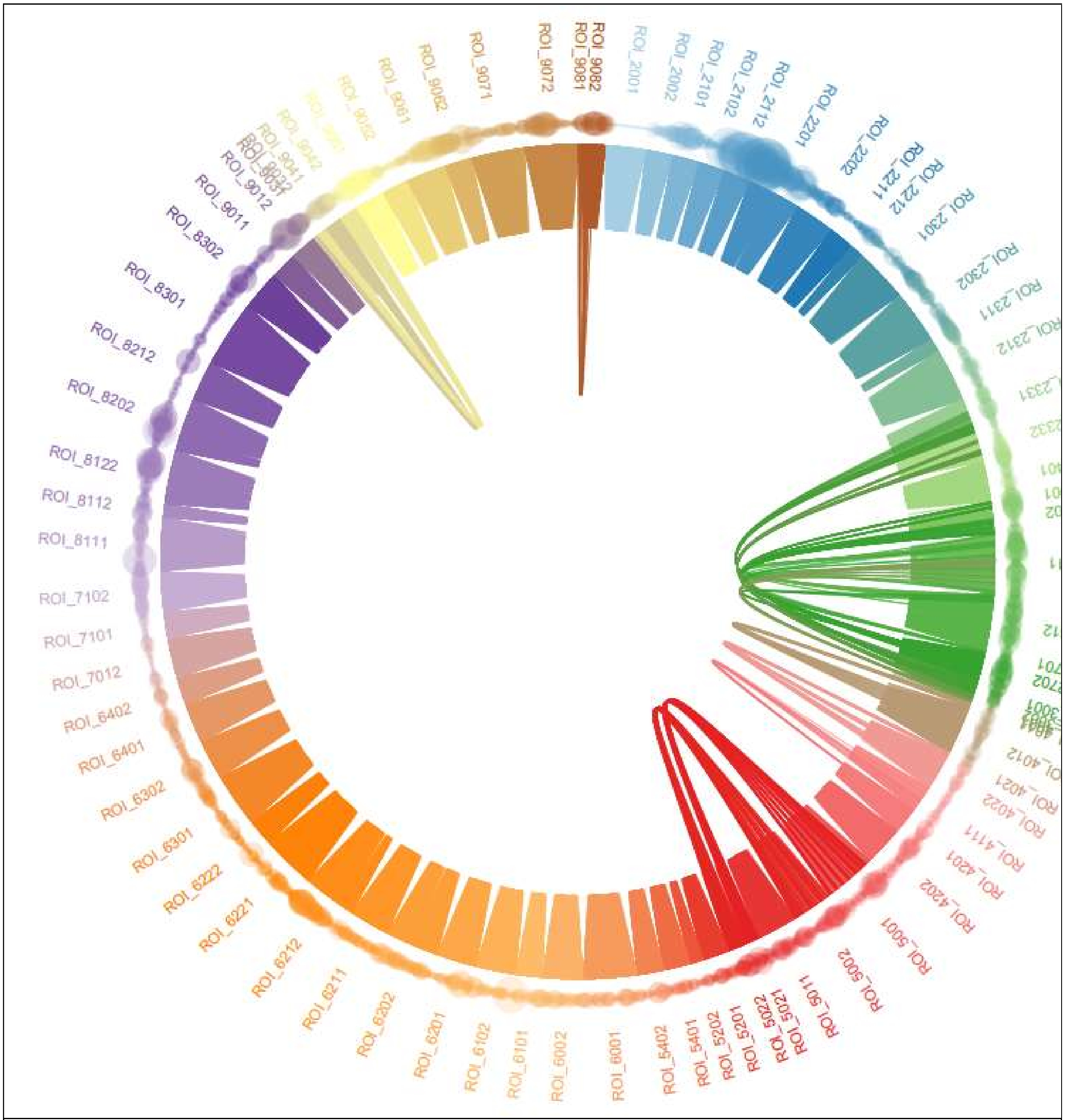


**Figure A2**: Local brain network connectome differentiating AD and healthy. Each dot represents a voxel and each cluster of arcs represents a local network detected. Different colors represent different ROIs. A network connecting dots in different colors indicates that the network is crossing multiple ROIs.

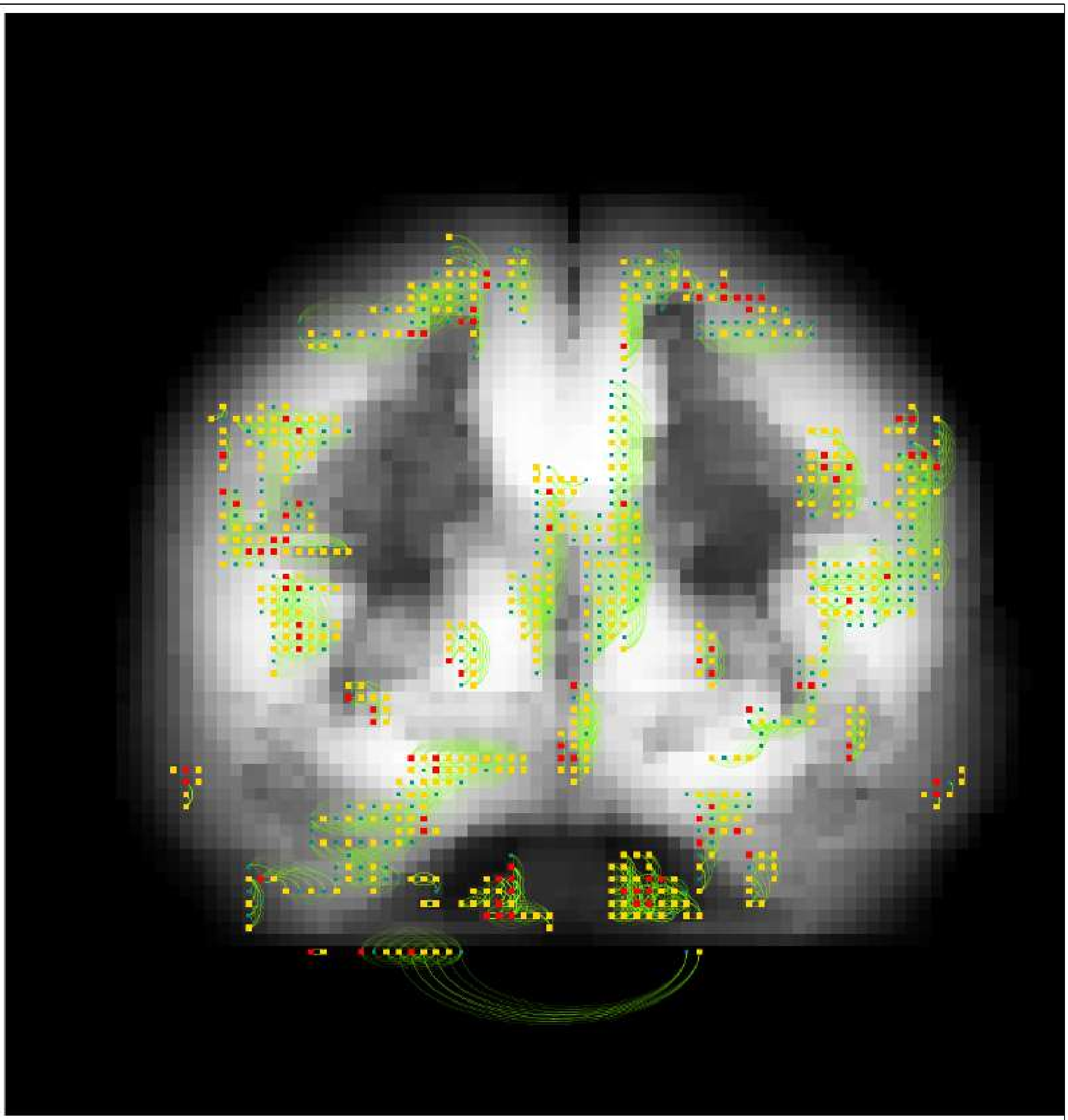

**Figure A3**: Overlaying of the brain network connectome differentiating AD and healthy. All local networks are projected to a coronal slice of the brain at midline. Red voxel: marginally strong signal. Yellow voxel: marginally weak signal. Blue voxel: non-selected connecting voxel in a selected local brain network (might differentiate other pair of classes).

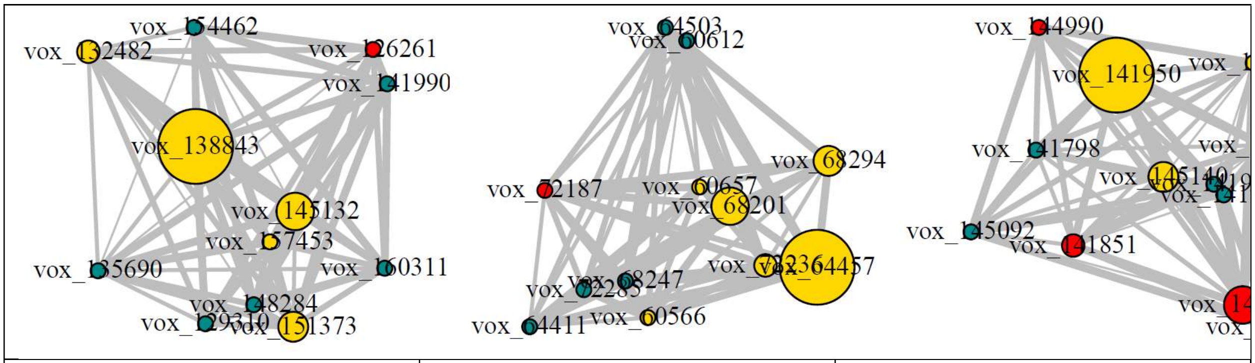

(a) vox_138843 & vox_145132 & vox_151373 & vox_132482
ROI: Precuneus_L

(b) vox_64457 & vox_68201
ROI: Frontal_Med_Orb_R

(c) vox_141950
ROI: Parietal_Inf_L

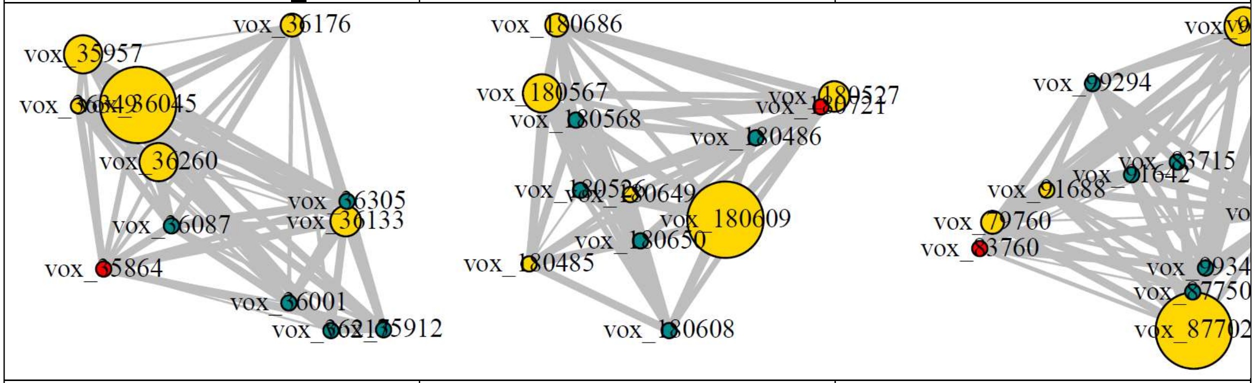

(d) vox_36045
ROI: Temporal_Mid_R

(e) vox_180609
ROI: Paracentral_Lobule_L

(f) vox_87702
ROI: Cingulum_Ant_L

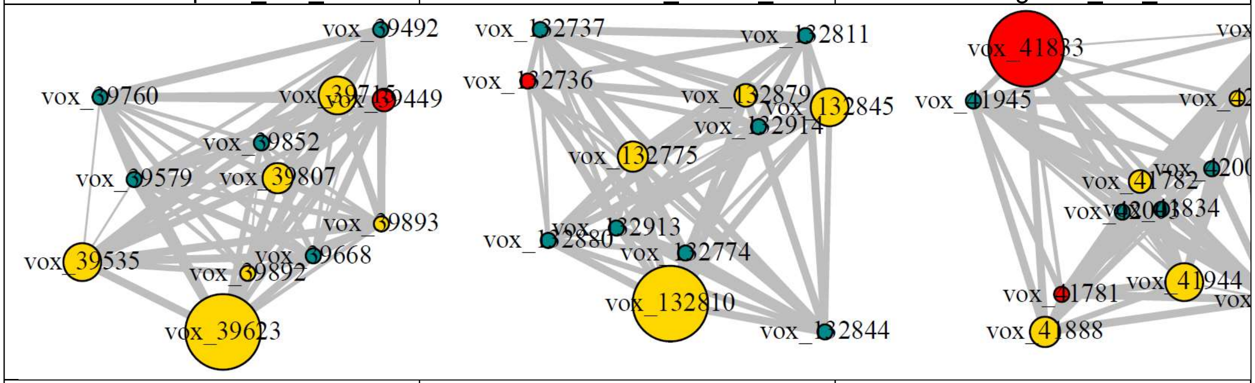

(g) vox_39623
ROI: Temporal_Mid_R

(h) vox_132810
ROIs: Cingulum_Mid_L & Cingulum_Mid_R

(i) vox_41833
ROI: Occipital_Inf_R

**Figure A4:** Example local brain networks that differentiate MCI and healthy groups. Yellow: marginally weak signals. Red: marginally strong signals. Blue: other selected connecting signals that differentiate other pairs. Node sizes are proportional to the local-network-adjusted mean differences. Edge widths are proportional to the absolute partial correlations. The top selected voxels within each network and the ROI(s) the network resided in are indexed.

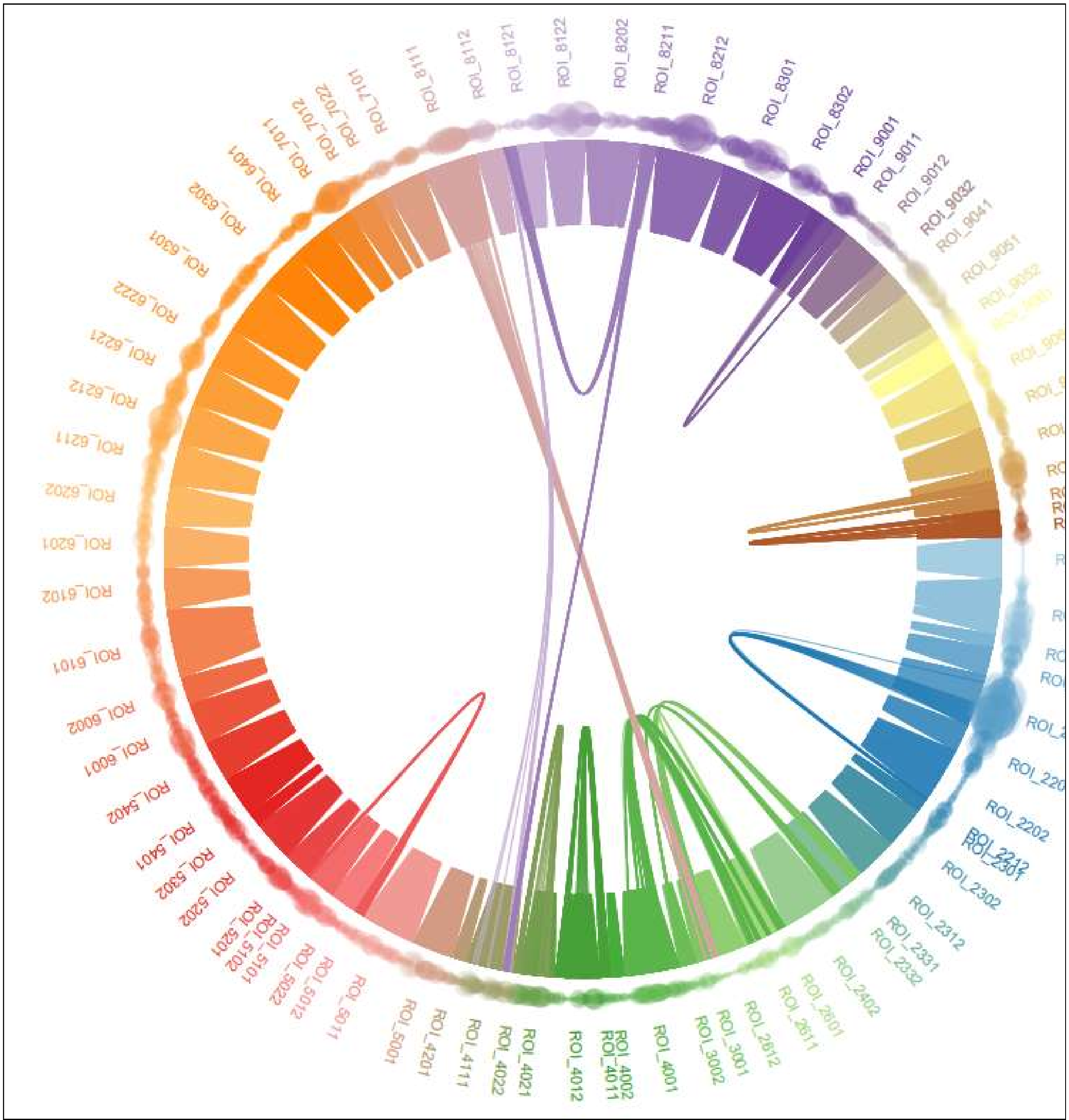


**Figure A5**: Local brain network connectome differentiating MCI and healthy. Each dot represents a voxel and each cluster of arcs represents a local network detected. Different colors represent different ROIs. A network connecting dots in different colors indicates that the network is crossing multiple ROIs.

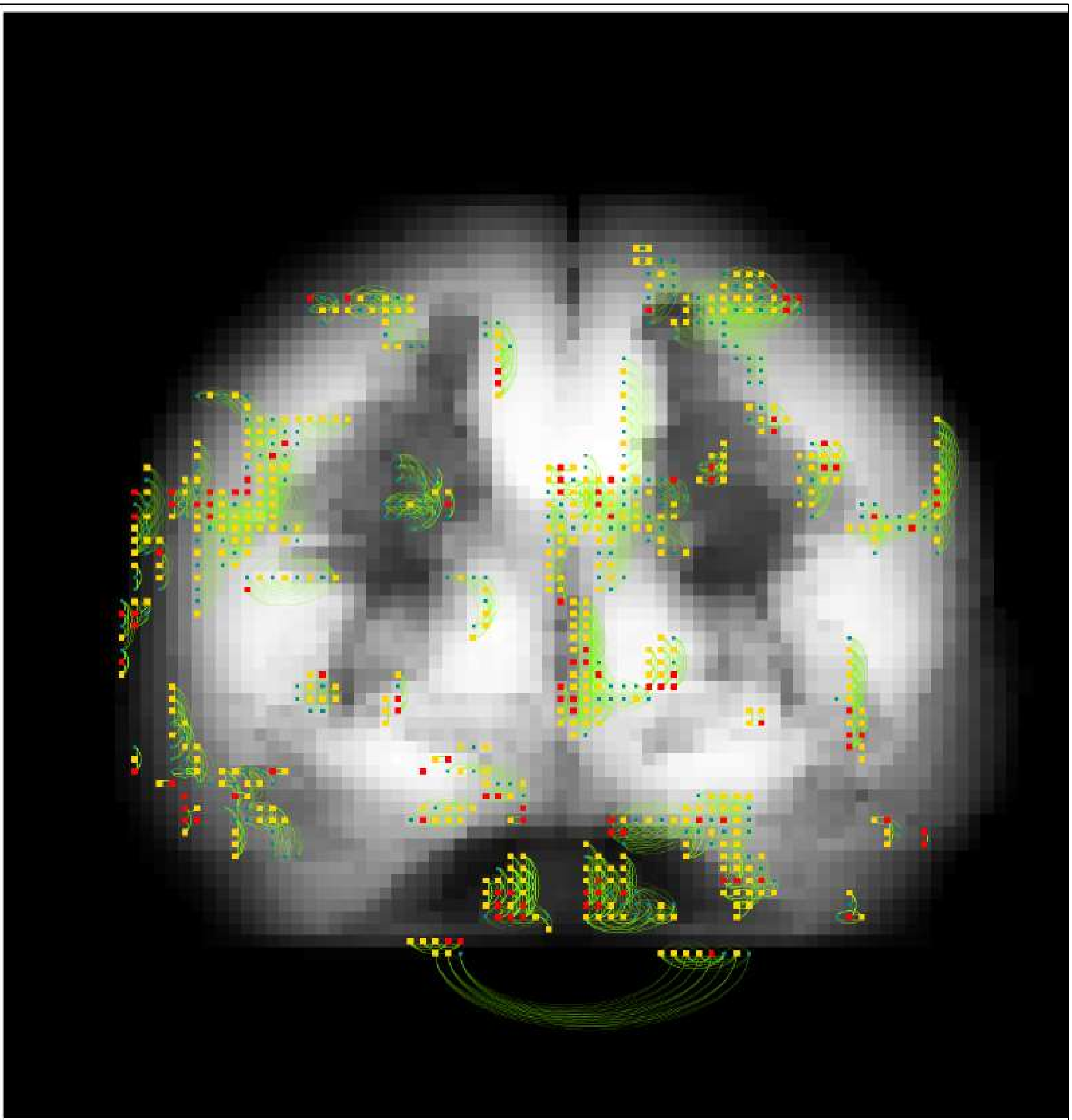

**Figure A6**: Overlaying of the brain network connectome differentiating AD and MCI on the brain. All local networks are projected to a coronal slice of the brain at midline. Red voxel: marginally strong signal. Yellow voxel: marginally weak signal. Blue voxel: non-selected connecting voxel in a selected local brain network (might differentiate other pair of classes).
